# Microbial degradation of plastic in aqueous solutions demonstrated by CO_2_ evolution and quantification

**DOI:** 10.1101/719476

**Authors:** Ruth-Sarah Rose, Katherine H. Richardson, Elmeri Johannes Latvanen, China A. Hanson, Marina Resmini, Ian A. Sanders

## Abstract

The environmental accumulation of plastics worldwide is a consequence of the durability of the material. Alternative polymers, marketed as biodegradable, present a potential solution to mitigate their ecological damage. However, understanding of biodegradability has been hindered by a lack of reproducible testing methods. Here, we present a new approach to assess biodegradability by monitoring bacterial respiration, in an aqueous media supplemented with a polymer as a single carbon source. *Rhodococcus rhodochrous* and *Alcanivorax borkumensis* are good model organisms for soil and marine systems respectively, allowing simulations of plastic biodegradation in these key environments. We demonstrate that polymer molecular weight reduction is the critical factor in the biodegradability of low-density polyethylene. Additionally, we tested a wide variety of plastics, including environmentally weathered and laboratory aged samples, using the same method allowing direct comparisons of the relative biodegradability of various polymers.

## Introduction

Plastic is a versatile material. It is waterproof, strong, easily shaped and can be made rigid or flexible. Furthermore, it is relatively cheap to manufacture, making it widely available across the world. However, the inherent stability and durability of this material has resulted in widespread accumulation of plastics in both terrestrial and aquatic environments^1^. Indeed, evidence of floating plastic debris can be found from the equator to polar ice caps^2^ and has led to the current global crisis^3–5^. There is considerable debate about possible solutions^6^. One key focus is the development and use of plastic alternatives, involving new degradable polymers such as hydrodegradable^7^, compostable^8^, biocomposites^9,10^, bioplastic^11^ and pro-oxidantadditive-containing (PAC)/oxo-degradable plastics^12^. Opinion is greatly divided over the environmental impact and the biodegradability of these materials^13^. This is largely due to the methods employed to measure biodegradation where biodegradation is commonly evidenced by either the loss of plastic mass or the monitoring of microbial respiration with both approaches using samples that have been buried in soil^14–16^. However, soil composition varies globally, thus the relevance and reproducibility of these tests have been questioned^8,14,17,18^. Additionally, these methods are not suitable for studying plastic decomposition in the open or aquatic environments, where oxygen and nutrient concentrations, temperature and microbial ecology differ significantly^17,19^. A number of alternative approaches have been reported in the literature^20–25^ but have not been widely applied. To this end, there is an urgent need for a relevant, reliable, versatile and standardised procedure for the measurement of biodegradability in aqueous conditions. This would allow comparative monitoring of the fate of different plastics and the evaluation of their environmental impact.

Here, we present a standardised and reproducible bioassay to quantitate the biodegradation of plastic in a defined minimal media, based on the detection and quantification of CO_2_, produced as a result of bacterial respiration, using gas-chromatography. We studied the biodegradability of different samples of low density polyethylene (LDPE), starch-based compostable and oxo-degradable plastic, using the soil bacterium *Rhodococcus rhodochrous*^20,26,27^. The PAC or oxo-biodegradable plastics are polyolefins blended with additives that stimulate the cleavage of polymer chains under oxidative conditions, a feature expected to lead to improved biodegradability^28^. Artificially aged and naturally weathered samples of were used to investigate how these parameters impacted biodegradability of each plastic. Sampling of LDPE and oxo-LDPE aged for different lengths of time allowed the study of the relationship between polymer molecular mass and degradation properties. The method was expanded to probe the activity of the marine bacterium *Alkanivorax borkumensis*^29,30^ on plastic to begin to explore degradation in seawater.

## Results and discussion

### Assessment of microbial growth

We initially assessed microbial growth on plastic, employing the soil bacterium *R. rhodochrous* as a model organism (Table S1) with previously reported methods^20–24^. Low density polyethylene (LDPE) was chosen as the target plastic, as it is one of the most common polymers used for packaging and dominates the composition of plastic waste on the sea surface^31^. Scanning electron microscopy (SEM), and SYBR green staining of *R. rhodochrous* grown on solid media with LDPE as the sole carbon source, showed microbial colonisation of the plastic surface within 20 days (Fig S1a). Efforts to dislodge cells from the film with multiple water washes, in order to estimate bacterial growth by cell counts, proved unsuccessful as bacteria were observed still attached to the film. More aggressive methods such as acid, detergent or bleach washes also failed to remove bacteria completely (Fig 1a and Fig S1). It is unlikely that this robust adhesion was specific to polymer type as there was no significant difference between the number of bacterial cells per mm^2^ when grown on either LDPE or oxo-LDPE (*p*=0.6, Fig S1d). This contradicts previous suggestions that additives, such as those found in oxo-LDPE, may alter the hydrophobicity of the polymer surface, thus influencing microbial attachment^27,32^. Biochemical assays supported the hypothesis of irreversible adhesion, as evidenced by a lack of measurable growth on LDPE in comparison to glucose (Fig S2). Consequently, any assay requiring the removal of biomass or a bacterial suspension would be fundamentally inaccurate^21,33^. It is noteworthy that substantial interference by samples of heat-treated plastic were observed, in the absence of any microbe, in both the luciferase (ATP/ADP) and AlamarBlue assays^34,35^.

**Figure 1.**
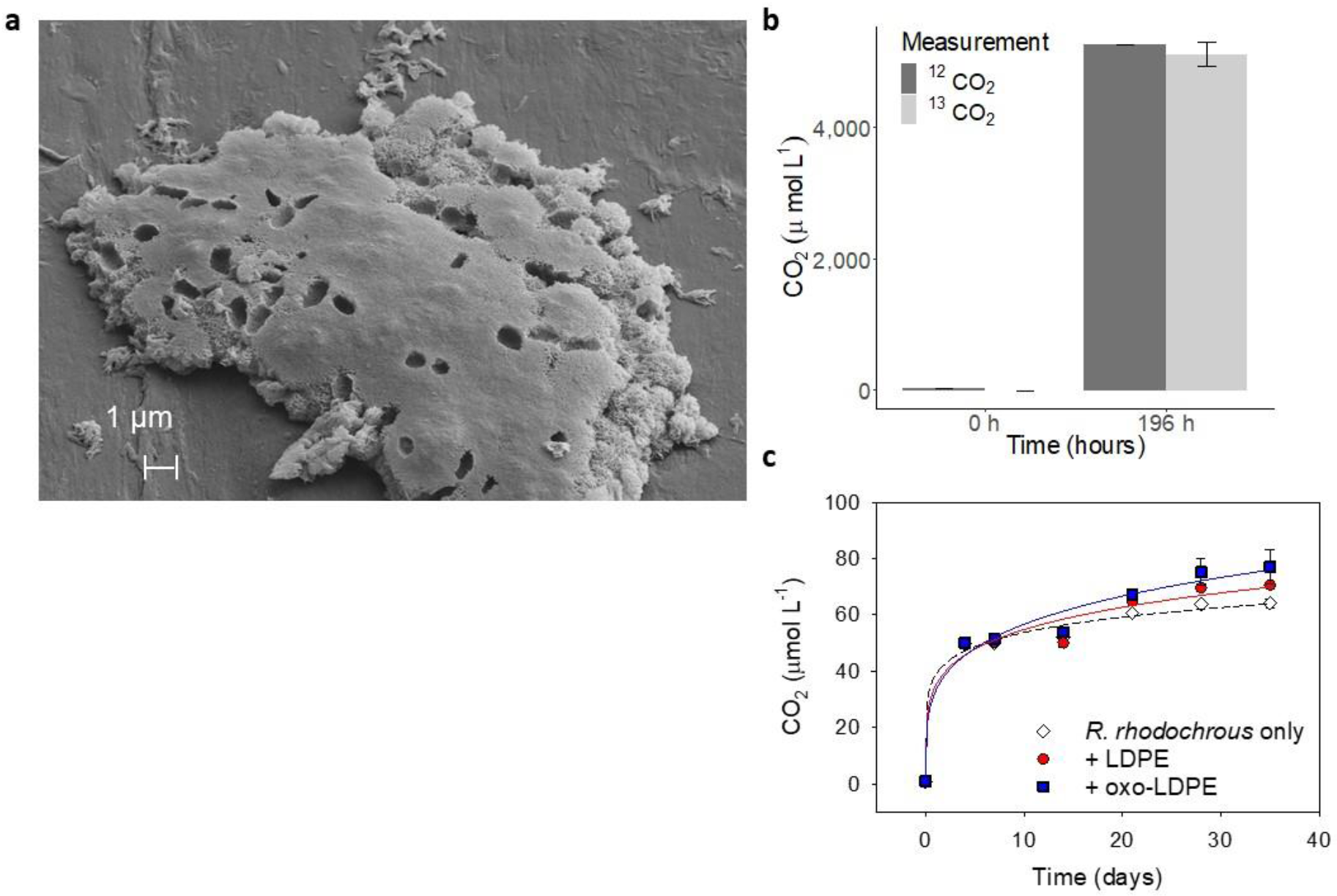
Microbial growth and measurement of biological degradation of polymer films. a) *R. rhodochrous* grown on LDPE underwent water washes and acid washing prior to visualisation by SEM (scale bar 1 μm). Large deposits of dead cells are clearly still visible after this treatment, revealing the challenges in the removal of bacterial films. b) *R. rhodochrous* grown on ^13^C-glucose. There is no difference between measurement of CO_2_ (dark grey) or ^13^CO_2_ (light grey) after 196 hours (*n*=3, *p*=0.98), directly linking carbon uptake with respiration. c) Measurement of CO_2_ production by *R. rhodochrous* in minimal media (open diamond), on unaged LDPE (red circle) or unaged oxo-LDPE (blue square) revealed no significant effect due to the presence or type of plastic (*n*=5, *p*=0.99).

### Measurement of growth by monitoring CO_2_ production

The use of gas-chromatography for the detection and quantification of CO_2_ produced by bacterial respiration has been used consistently in environmental science^36,37^. However, the high throughput GC approach has not been explored as a tool for the evaluation of polymer biodegradation. We modified the method to detect CO_2_ evolved from a well-defined aqueous media containing a single microbial culture and a single type of polymer, as the only carbon source. Thus, the produced CO_2_ can be directly linked to the mineralisation of the supplied carbon source via bacterial respiration. To confirm the validity of the assay, growth of the soil bacterium *R. rhodochrous* was carried out on ^13^C-glucose^38^. The data (Fig 1b) show that the concentration of CO_2_ after 196 hours was consistent with that produced as ^13^CO_2_, confirming the suitability of our methodology (*p*=0.98). At the point at which stationary phase of growth was achieved, 32 % of the available carbon had been released as CO_2_, with the majority of carbon likely assimilated into biomass^39,40^.

The detection of evolved CO_2_ to monitor respiration, and thus growth, of *R. rhodochrous* on samples of unaged LDPE and oxo-LDPE, was carried out over 35 days (Fig 1c). Within the initial days, ~0.05 mM CO_2_ was released, from all cultures, most likely due to carry over from the starter culture that was saturated with highly soluble CO_2_. Thereafter, stationary phase was achieved, with no significant difference between *R. rhodochrous* grown with no carbon source and the bacteria grown on unaged LDPE or oxo-LDPE (*p*=1). Sample acidification, from the release of CO_2_, was ruled out as an explanation for the limited growth as the pH of the media remained unchanged during the incubation. The available concentration of phosphate (24 mM) and nitrogen (5.6 mM) in the starting media were too high to limit the production of 0.05 mM CO_2_. Thus, the data suggest that bio-available carbon was the limiting factor in microbial growth and that unaged LDPE and oxo-LDPE show limited evidence of biodegradation.

### Effect of UV aging on biodegradation

Polymers found in the environment are subject to chemical changes as a result of heat, light and water^41^. Artificial aging by sustained UV exposure is commonly used to simulate environmental damage as it allows reproducible samples to be generated^42^. We investigated the role of artificial UV aging in the biodegradability of LDPE and oxo-LDPE using samples of films irradiated for 450, 758 and 900 hours, in accordance with international standards (ASTM 5208 Cycle C). The films, unaged and aged, were incubated with *R. rhodochrous* for 35 days and CO_2_ measured over time. As expected, a clear link between UV aging and biodegradation was observed, which was further enhanced in the presence of the oxo-additive. Indeed, the biodegradation of oxo-LDPE (+450 h UV) was 90-fold greater than LDPE (+450 h UV) and 45-fold greater than unaged oxo-LDPE after 35 days (Fig 2a). However, biodegradation of LDPE (+450 h UV) was only 3-fold greater than unaged LDPE (*p*=0.26) demonstrating the impact of the additive. Surprisingly, longer UV exposure of oxo-LDPE samples, for 758 and 900 hours, did not increase CO_2_ evolution beyond that observed after 450 hours. Indeed, less CO_2_ was released, though there was no significant difference between CO_2_ evolved for oxo-LDPE (+758 h UV) and oxo-LDPE (+900 h UV) (*p*=0.67) (Fig S3). The reasons for this are not fully understood and require further investigation. Conversely, biodegradation of LDPE continued to increase with irradiation time, such that the samples of both LDPE and oxo-LDPE that had undergone 900 hrs of exposure generated similar quantities of CO_2_ (*p*=0.59). This demonstrates that lengthier UV exposure is necessary to facilitate biodegradation of conventional LDPE while, in contrast, it impairs the biodegradation of oxo-LDPE.

**FIGURE 2.**
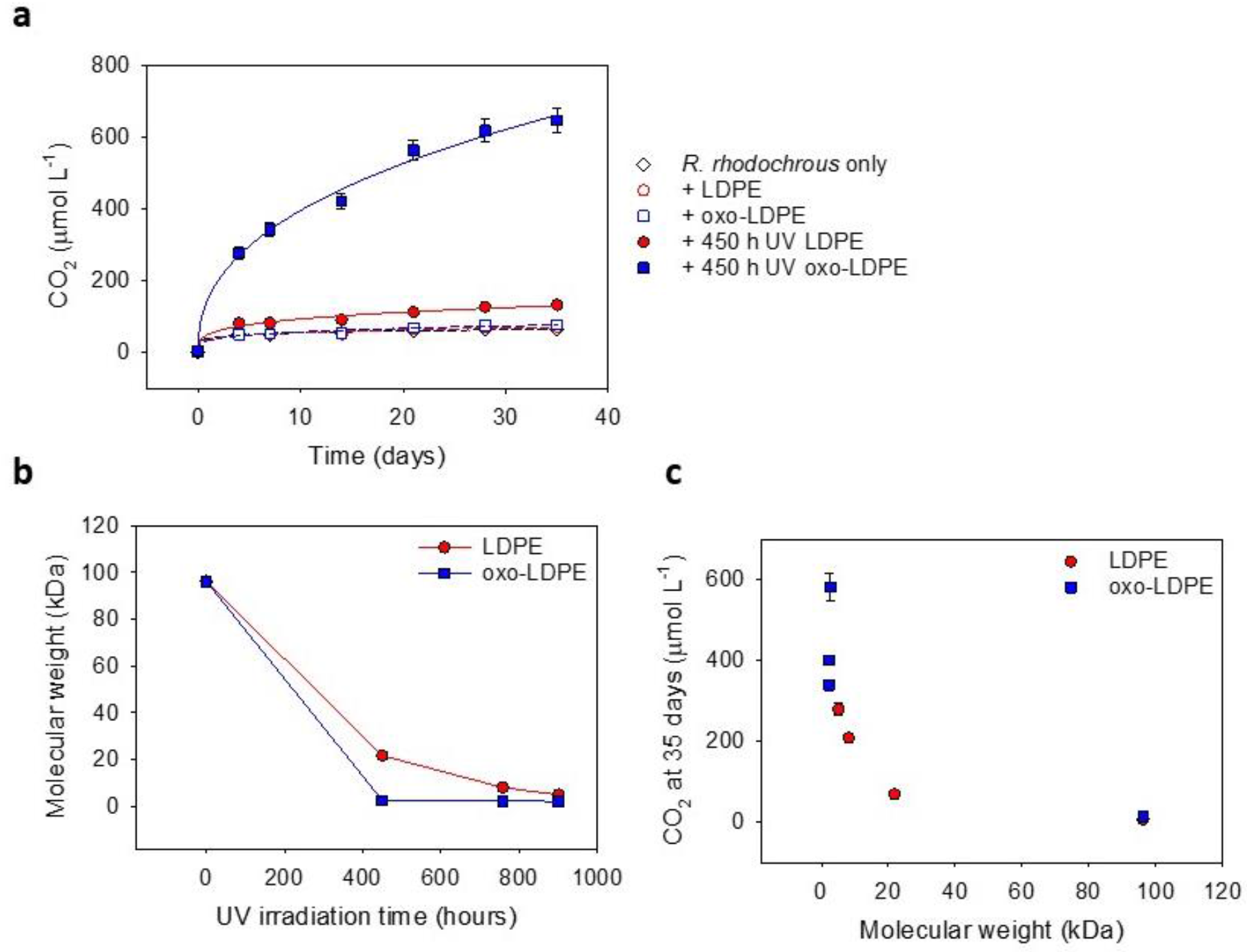
Artificial UV ageing accelerates the rate of biodegradation of oxo-LDPE compared to LDPE as a result of changes in molecular weight. a) CO_2_ production from *R. rhodochrous* (*n*=5) grown with no supplied carbon (open black diamond), on unaged LDPE (open red circle), unaged oxo-LDPE (open blue square) compared to 450 hours UV irradiated plastic of both LDPE (solid red circles) or oxo-LDPE (solid blue squares). UV exposure of oxo-LDPE significantly affects CO_2_ production (*p*<0.001) while there is only a small increase in CO_2_ production between UV treated LDPE and unaged plastics (*p*=0.26). b) The molecular weight (kDa) of LDPE (red open circles) and oxo-LDPE (blue solid squares), as measured by gel permeation chromatography, compared to UV exposure time (*n*=3). c) The concentration of CO_2_ measured at 35 days for *R. rhodochrous* grown on LDPE (red) or oxo-LDPE (blue) after UV irradiation (*n*=5, error bars represent standard error) compared to the molecular weight of the sample reveals a clear link between molecular weight and CO_2_ production.

### Influence of molecular weight on biodegradation

UV irradiation of both LDPE and oxo-LDPE polymers is known to result in changes in the chemical structure of the matrix, as a result of oxidation^12,43^. To understand the influence of UV-ageing on biodegradation, changes in polymer molecular weight as a function of irradiation were measured (Fig 2b). Here, there is a marked difference in the time required for the polymer samples to reach a molecular mass <3 kDa. Oxo-LDPE achieves this reduction within 450 hours of irradiation, whereas LDPE requires double the exposure time (900 hours) to reach the same mass. When the different samples were incubated for 35 days with *R. rhodochrous* and CO_2_ measured, a direct correlation between CO_2_ produced and molecular mass of the polymer sample was observed (Fig 2c). The greatest concentration of CO_2_ was generated by microbial action on oxo-LDPE after 450 hours, when the molecular mass reached <3 kDa. This is consistent with the theoretical proposal that only polymers of less than 5 kDa are bioavailable^44^. The data appear to suggest that under the experimental conditions used, oxo-LDPE can be considered more biodegradable than conventional LDPE, as a result of reaching a lower molecular mass in a shorter period of time. The experiments allow comparison between different types of polymers artificially aged, although these findings would need to be validated with naturally aged samples.

### Starch based plastic

“Compostable” plastics are often derived from biomass^45^. Here, the polymers contain carbonyl or ester functional groups within the backbone making them particularly prone to hydrolysis^25,46^. These plastics are designed to biodegrade under composting conditions, *i.e*. a soil-based, warm, anaerobic environment. *R. rhodochrous* is a bacterium found in soil and may play a role in the decomposition of plastic in the natural environment; thus, a sample of commercially available starch-based compostable plastic was also assayed (Fig S4, Table S4). The data show that *R. rhodochrous* released 2.5 times more CO_2_ when grown on a compostable plastic than on LDPE but 5.5 times less than on oxo-LDPE after 450 hours of UV irradiation. The ability to utilise compostable plastic as a substrate is likely related to the composition of the polymer and is consistent with more recent findings that suggest compostable plastics may be degraded more readily than all other forms of plastic^25^.

### Biodegradation in water

Artificial aging of plastic is routinely applied and accepted practice (ASTM 5208), as it allows accelerated decomposition and the preparation of reproducible samples. However, it is not entirely representative of environmental weathering^41,47,48^. To this end, prior to testing, samples of LDPE and oxo-LDPE were surface-weathered in sea water for 82 days, undergoing natural variations in sunlight and UV intensity. Scanning electron microscopy (SEM) revealed evidence of damage on both polymer films due to microbial activity (Fig 3a) and the presence of microorganisms still attached to the film (Fig 3c-d). Additionally, extensive calcium phosphate deposits were identified (Fig 3b); these could dissolve during incubation, altering the media composition and influencing bacterial growth. Consequently, it was difficult to ensure consistent sampling of environmentally weathered samples. Until a method that allows complete removal of bacteria without altering the polymer, as is the case with autoclaving/heating, a large sample size is going to be critical when assessing the biodegradation of environmental samples.

**FIGURE 3.**
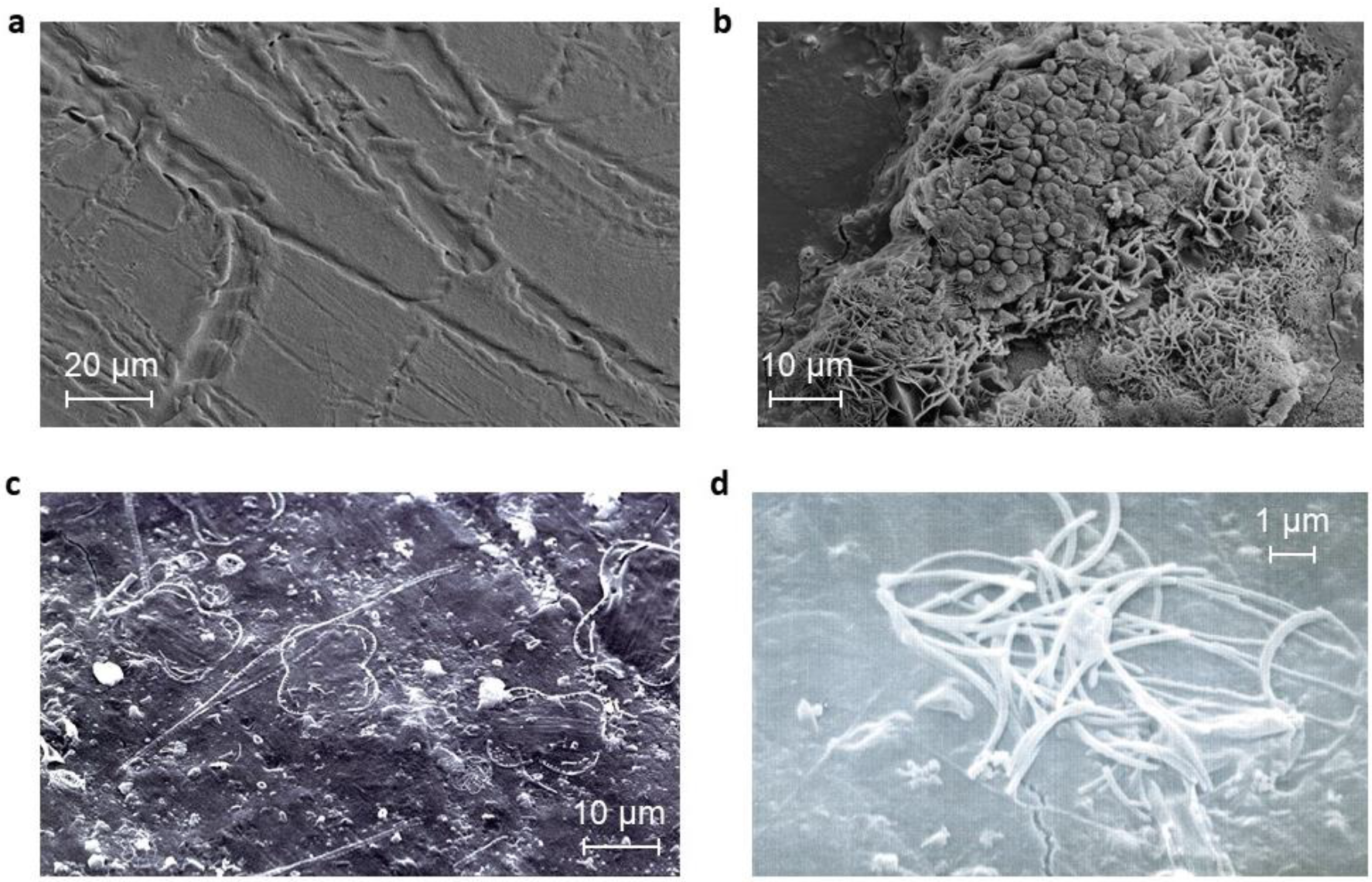
Microbial growth on plastic in environmental samples may influence testing. Oxo-LDPE was suspended on the surface in sea water tanks for 82 days. The plastic was washed and visualised by SEM. a) tracks consistent with hyphael damage were observed (error bar 20 μm). b) Salt deposits were observed (error bar 10 μm). c) Evidence of crystallised microbes on the plastic surface (error bar 10 μm). d) Residual microbial presence was identified (error bar 1 μm).

There is increased interest in the microbial composition of aquatic environments and, more specifically, microbes that play a role in the degradation of plastic in the sea^49^. To this end, we used our bioassay with the model organism *Alcanivorax borkumensis* for studying hydrocarbon metabolism within the marine environment^50^. The growth media was altered to accommodate the higher salt concentration required to cultivate *A. borkumensis* since it was unable grow in the same media used for *R. rhodochrous*. Incubation of *A. borkumensis* with pyruvate as a natural carbon substrate generated a typical growth curve (Fig. 4a). Stationary phase was achieved after 10 days by when *A. borkumensis* had respired 26 % of the available carbon (Fig 4a).

**FIGURE 4.**
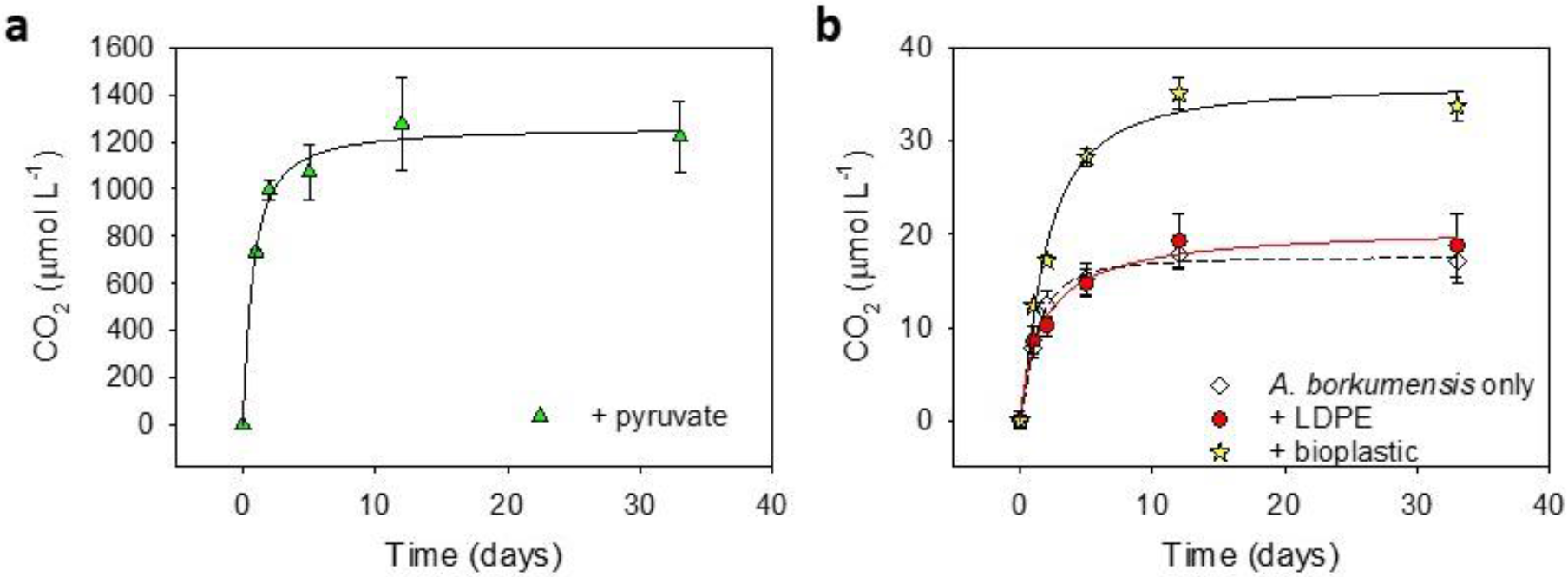
Monitoring respiration of *A. borkumensis* when grown on a variety of substrates. a) *A. borkumensis* was grown in liquid culture with pyruvate as the sole carbon source and CO_2_ measured over time (*n*=4, error bars represent standard error). b) *A. borkumensis* was grown in liquid culture with no carbon source (open diamonds), LDPE (red circles) or with a bioplastic (yellow star) and CO_2_ production was measured over time by GC. *A. borkumensis* evolved twice as much carbon when grown on bioplastic compared to bacteria or LDPE alone (*n*=5, *p*=0.0006).

### Monitoring of *A. borkumensis* growth on plastics

*A. borkumensis* has been shown to be capable of growth on larger alkanes of up to C38^50^; however, its ability to metabolise polyethylene has not been investigated., *A. borkumensis* was grown on LDPE and CO_2_ production monitored. Here, as for *R. rhodochrous*, there was an initial release of CO_2_ prior to a plateau in CO_2_ production. There was no significant difference between CO_2_ generated by *A. borkumensis* on LDPE and the sample without plastic (*p*=0.88) (Fig 4b), revealing that *A. borkumensis* is unable to utilise LDPE as a substrate.

“Bioplastics” can refer to polymer composites^14^ and are marketed as a biodegradable alternative to LDPE. *A. borkumensis* was grown in liquid culture with a polymer composite, and CO_2_ production monitored over time. Interestingly, there was only a two-fold increase in CO_2_ evolved from *A. borkumensis* in the presence of the polymer film compared to *A. borkumensis* alone (*p*=0.0006) (Fig 4b). Over the incubation, insufficient CO_2_ was released to suggest that any nutrient, other than carbon, had become limiting. The bioplastic was more biodegradable than LDPE but 40 times less than pyruvate.

## Conclusions

To date, assessing the biodegradability of plastic has relied upon either composting approaches i.e., burying samples in soil, which have shown limitations in reproducibility^14,15^ or samples being suspended in the sea, where samples risk being lost prior to recovery for quantification^25^. Our method allows the reliable assessment of plastic biodegradability via the quantification of CO_2_ produced in aqueous media as a direct result of microbial metabolism. By bio-assaying *Rhodococcus rhodochrous*, from soil, and *Alkanivorax borkumensis*, from sea water we have demonstrated that the choice of bacterial strain influences polymer degradation. The method is reliable, reproducible and allows microbial activity to be monitored either without interfering with the cell culture or trying to remove adhered cells, reducing the risk of contamination or disrupting biofilms. The method was adapted to accommodate the different nutrient requirements for each species, thus could be adapted to imitate a specific aquatic environment. As the microbiological ecosystems of terrestrial, freshwater and marine habitats are inherently different, this approach will allow the assessment of the role of individual bacterial strains or communities in plastic biodegradation.

The key role that abiotic degradation plays in the biological decomposition of plastic is highlighted by our experiments on naturally weathered or artificially aged samples of LDPE and oxo-LDPE. The method presented here, allowed the measurement of biodegradability irrespective of the chemistry of the polymer. Our data demonstrate a dependence between biodegradability and molecular mass of plastic samples. The aged oxo-LDPE samples form fragments of lower mass in a shorter period of time and this results in a significantly higher amount of CO_2_ detected, therefore indicating a direct relation between the rate of biodegradability and the speed at which the molecular mass of the polymer is reduced. The wider applicability of the method is demonstrated by testing compostable plastic and bioplastics and showing that these are more readily degraded than unnamed LDPE samples. This is the first example of work where ageing, chemical structure of plastic and biodegradability have been connected. The method provides a robust and reproducible approach for comparing different types of polymers and evaluate the effect of environmental and/or artificial ageing.

## Materials and Methods

### Cell Culture

*Rhodococcus rhodochorous* (ATCC-29672 also termed *Rhodococcus ruber*) and *Alcanivorax borkumensis* (ATCC-700651) were cultured in media as recommended. Each bacterium was grown over night in minimal media (Table S2 and S3). *R. rhodochorous* was grown at 27 °C, shaking at 120 rpm and *A. borkumensis* was grown at 30 °C, shaking at 120 rpm. Bacteria were handled in a class II biosafety cabinet throughout.

### Plastic preparation

Full details of all the plastic used her can be found in Table S4. Plastic was washed with 70 % ethanol and allowed to dry overnight in a microbiological safety cabinet prior to use in each assay.

### Preparation of bacteria on minimal media plates

Plates containing minimal media with no carbon source were prepared. Bacterial suspensions in the exponential phase were prepared and a concentration of 1×10^8^ cells ml^−1^ were applied to the plates. Four sterile plastic pieces of 1cm^2^ were placed on top of the cultures. The plates were allowed to dry before sealing with parafilm, inversion and incubation at 30 °C.

### Scanning Electron Microscopy

Plastic samples in liquid medium were rinsed briefly in deionised water, air dried and placed on double sided self-adhesive carbon discs on aluminium stubs. Plastic samples grown with bacteria on agar plates were carefully lifted from the plate and placed directly on double sided self-adhesive carbon discs on aluminium stubs. The mounted samples were coated with approximately 7.5 nm of gold in a Cressington 108 auto sputter coater, the thickness being controlled with a Cressington MTM 10 thickness monitor. Coated samples were examined and imaged using secondary electrons in either a Zeiss Sigma Gemini scanning electron microscope, with a field emission electron gun at an acceleration voltage of 10 kV, or in a JEOL JSM 6480LV scanning electron microscope at an acceleration voltage of 30 kV.

### Preparation of bacteria for the plastic CO_2_ bioassay

A starter culture was prepared in liquid media and grown over night at 30 °C. To separate the bacterial cells, sterile glass beads were added to the overnight culture and mixed. The solution was diluted 1:4 in minimal media and the cell number estimated by measuring the optical density at 600 nm, as per standard methodology. To estimate the cell number, the relationship between optical density and cells number for *E. coli* was used (OD_600_= 1.0= 8 x 10^8^ cells ml^−1^). The inoculum was thoroughly mixed and diluted in minimal media to achieve 7.8 x 10^8^ cells ml^−1^. A 0.4 ml volume was applied to each autoclaved container, 0.5 mg of sterile plastic added where required, the tubes sealed and the cultures incubated at 27 °C, shaking at 120 rpm. Five replicates of each test condition were prepared in 12 ml gas-tight vials (Exetainers, Labco, Ceredigion, UK) for measurement of CO_2_ production.

### CO_2_ analysis

At various time points throughout the assays, the samples were removed from the incubator for GC analysis and returned to the incubator as soon as possible. A 50 μl sample was withdrawn from the headspace using a gas tight syringe in an auto-sampler (Multipurpose Sampler MSP2, Gerstel, GmbH, Germany) and injected into a gas chromatograph fitted with a flame ionising detector (GC-FID) and hot-nickel catalyst (Agilent Technologies UK Limited, Cheshire) to reduce the CO_2_ to CH4. Conditions were: column at 30 °C, detector at 375 °C and catalyst at 385 °C. CO_2_ was separated using a stainless-steel column (length 6’ x ½ø) packed with Porapak (Q 80/100), and with a hydrogen/air mix (7/93%, Zero grade BOC) as the carrier gas (430 ml min^−1^). Headspace concentrations of CO_2_ as CH4 were calculated from peak areas using an electronic integrator. The concentration of CO_2_ was calibrated against a known standard (3700 ppm) (BOC, UK) and air.

### ^13^C glucose analysis

Samples (as above) were incubated in 3.5 ml gas-tight vials (Exetainers, Labco, Ceredigion, UK). Gas samples, 30 μl, were taken from the headspace of the ^13^C glucose and control incubation vials and injected into 3.5 ml exetainers. These were pre-injected with CO_2_ to act as a carrier for the sample giving a final concentration of 1100 ppm. A 500 μl gas sample was withdrawn from the headspace using a gas tight syringe in an auto-sampler (Multipurpose Sampler MSP2, Gerstel, GmbH, Germany) and injected into a flash elemental analyser (1112 Series, Thermo, Bremen, Germany), interfaced with a continuous flow isotope ratio mass spectrometer (Sercon 20/22, Crewe, UK). CO_2_ was separated from other gases on a GC Poropak column (PoraPLOT Q). Carbon isotope calibration utilised the international standard for carbon, Ref. 8542, sucrose, −10.47 ‰ δ13C *versus* Vienna-PeeDee Belemnite (VPDB), National institutes of Standards and Technology.

### Data fitting

There was a significant difference between the concentration of CO_2_ released from the media over the course of the experiment (*p*<0.01). Therefore, averages of CO_2_ produced at each time point for the media alone or media containing the relevant plastic were subtracted as a baseline. ANOVA and post-hoc Tuckey tests were performed using the statistical package R. Statistics were performed on all data points and the final time point.

The authors considered the application of the Monod equation to quantify respiration rates^39^. However, it was not possible, in this instance, to accurately define the concentration of carbon substrate due to distribution of macromolecular weight.

### Gel permeation chromatography

GPC was performed by Smithers Rapra and Smithers Pira Ltd (UK).

### ATP assay

Three replicates of each test condition were prepared in 3 ml vials. The reactions were incubated at 27 °C, shaking at 120 rpm. At each time point, 0.2 ml of extractant B/S was added to the sample and mixed well. The ATP assay was carried out using the BioThema ATP Biomass kit (BioThema, Sweden), following the manufacturer’s instructions, using a multiwell plate reader (BMG LABTECH, GmbHA).

## Acknowledgments

We acknowledge the skill and expertise of Dr R Angold for the SEM. We thank Prof M. Trimmer for expertise regarding GC. We acknowledge Birgit Solheim Huseklepp and Zara Rehman Malik for their contribution to supplementary figure 1 and Dr Chloe Economou for technical support. RR is grateful to Dr ERG Main and Mr S Wilson for stimulating discussions and advice.

## Author Contributions

R.R., K.R, E.J.L and I.S. designed and carried out the experiments. All analysed data. R.R, C.A.H. and M.R. prepared the manuscript.

