## Supplementary for "Microbial degradation of plastic in aqueous solutions demonstrated by CO_2_ evolution and quantification"

**Table S1. Advantages and disadvantages of reported methods for the either the biochemical or cell staining approaches to quantify microbial growth on plastic.**

**Biochemical and spectroscopic methods:**

| Method | Reference | Advantages | Disadvantages |
|---|---|---|---|
| ATP/ADP | [1] | High throughput; good for detecting cell viability | Cells must be detached from polymer; Potential polymer interference |
| Optical density at 600nm | [2] | High throughput | Measurement of turbidity which may be influenced by microplastics; cells must be detached from plastic and in suspension |
| Resazurin (AlamarBlue®) | [3,4] | High throughput; good for detecting cell viability and integrity | Chemistry of degradation may affect the assay; cells must be detached from plastic and in suspension |

**Cell counting approaches:**

| Method | Reference | Advantages | Disadvantages |
|---|---|---|---|
| Colony forming units | [5] | Detect viable cells only; can be useful if culturing a variety of cells on the plastic | Must detach cells from plastic; consumable heavy; slow process |
| Live/dead assay | [5] | Good to distinguish live cells and for quantification | Lengthy, technical; requires separate live and dead internal technical controls |

6. Fuller, W. A. *Introduction to statistical time series*. (Wiley, 1996).

**Table S2. A complete description of the media preparation for *R. rhodochrous.*** Liquid media was prepared fresh by adding 50 mL of additive to 950 mL minimal media to make up to 1 L.

**Minimal media:**

| **Chemical** | **Mass** |
|---|---|
| di-sodium phosphate dodecahydrate anhydrous ($Na_2HPO_4$) | 1.5 g |
| Potassium di-hydrogen phosphate anhydrous ($KH_2PO_4$) | 1.8 g |
| Sodium chloride (NaCl) | 0.5 g |

Dissolved to a final volume of 950 mL in $dH_2O$ (15 MΩ) and sterilised at 121 °C for 15 minutes.

**Additive:**

| **Chemical** | **Mass** |
|---|---|
| Magnesium sulfate heptahydrate ($MgSO_4.7H_2O$) | 0.02 g |
| Ammonium iron (II) sulfate hexahydrate ($(NH_4)_2Fe(SO_4)_2.6H_2O$) | 0.03 g |
| Calcium chloride hexahydrate ($CaCl_2.6H_2O$) | 0.015 g |
| Ammonium chloride ($NH_4Cl$) | 0.3 g |
| Trace elements | 10 µl |

Dissolved to a final volume of 50 mL $dH_2O$ (15 MΩ) and filter sterilised through a 0.2 µm filter.

**Trace element solution:**

| **Chemical** | **Mass** |
|---|---|
| Manganese sulfate tetrahydrate ($MnSO_4.4H_2O$) | 0.059 g |
| Boric acid ($H_3BO_3$) | 0.029 g |
| Zinc sulfate heptahydrate ($ZnSO_4.7H_2O$) | 0.022 g |
| Sodium molybdate dihydrate ($Na_2MoO_4.2H_2O$) | 2.35 g |
| Cobalt nitrate ($Co(NO_3)_2$) | 0.02 mg |
| Copper sulfate ($CuSO_4$) | 0.02 mg |

Dissolved to a final volume of 500 mL $dH_2O$ (15 MΩ), filter sterilised through a 0.2 µm filter and stored at 4 °C.

**Table S3: A complete description of the media (ONR7A) preparation for *A. bokumensis.*** The solutions were prepared and autoclaved separately and were combined once cooled to prepare 1 L of culture media.

**Solution 1:**

| **Chemical** | **Mass** |
|---|---|
| Sodium chloride (NaCl) | 22.79 g |
| Sodium sulfate ($Na_2SO_4$) | 3.98 g |
| Potassium chloride (KCl) | 0.72 g |
| Sodium bromide (NaBr) | 83.00 mg |
| Sodium carbonate ($NaHCO_3$) | 31.00 mg |
| Boric acid ($H_3BO_3$) | 27.00 mg |
| Sodium fluoride (NaF) | 2.60 mg |
| Ammonium chloride ($NH_4Cl$) | 0.27 g |
| di-sodium phosphate dodecahydrate dihydrate ($Na_2HPO_4.2H_2O$) | 47.00 mg |
| TAPSO | 1.30 g |

The powders were dissolved in dd$H_2O$ (18.2 MΩ), the pH adjusted to pH 7.6 with NaOH and a final volume of 500 mL achieved,

**Solution 2:**

| **Chemical** | **Mass (g)** |
|---|---|
| Magnesium chloride hexahydrate ($MgCl_2.6H_2O$) | 11.18 g |
| Calcium chloride dihydrate ($CaCl_2.2H_2O$) | 1.46 g |
| Strontium chloride hexahydrate ($SrCl_2.6H_2O$) | 24.00 mg |

Dissolved to a final volume of 450 mL in dd$H_2O$ (18.2 MΩ).

**Solution 3:**

| **Chemical** | **Mass (g)** |
|---|---|
| Ferrous chloride tetrahydrate ($FeCl_2.4H_2O$) | 2.00 mg |

Dissolved to a final volume of 50 mL in dd$H_2O$ (18.2 MΩ).

**Table S4. A description of the plastics used in the study**

| **Plastic** | **Polymer** |
| --- | --- |
| LDPE | 100% low density polyethylene |
| Oxo-LDPE | low density polyethylene + 1 % DG12-08; Symphony Environmental Ltd. |
| Compostable | Starch based compostable; off the shelf |
| Bioplastic | Film, compostable duplex laminate |

**Figure S1. Visualisation of microbial growth on plastic film.** a) *R. rhodochrous* was grown on solid media with LDPE as the sole carbon source for 14 days. The film was washed with water and stained with SYBR green (scale bar = 9 µm). b) SEM image of *R. rhodochrous* grown on oxo-LDPE. The film was subjected to bleach wash prior to imaging; the arrow indicates the presence of remaining bacteria (scale bar = 10 µm). c) SEM image of bacteria grown on oxo-LDPE where the film was acid washed before imaging (scale bar = 200 nm). d) *R. rhodochrous* was grown on solid media with LDPE (red) or oxo-LDPE (blue) for 14 days. The film was removed and repeatedly washed with water to remove unbound or dead cells. The cells were stained with SYBR green and counted ($n$=3). There was no statistical significance between colonisation of the two plastics ($p$=0.6).

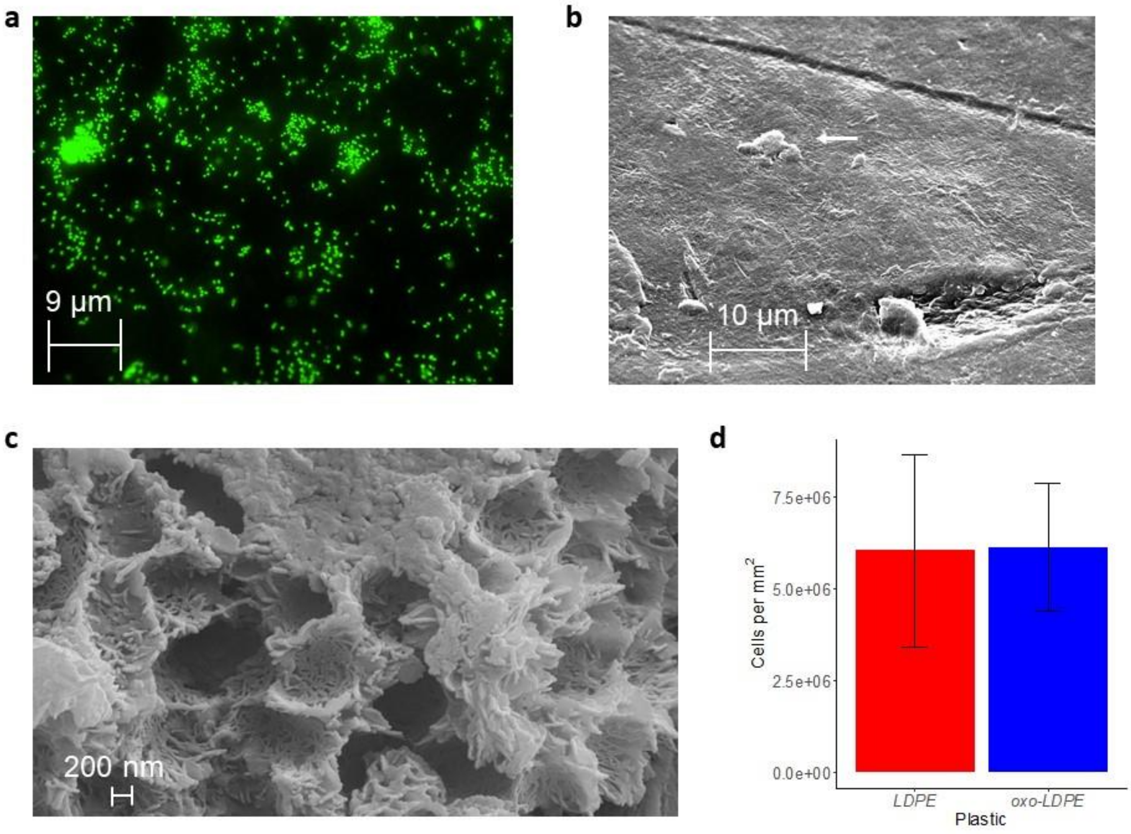

**Figure S2. Biochemical methods to measure microbial growth.**
a) The rate of ATP production measured using the ATP/ADP approach from cells grown in suspension with glucose (cyan triangles) as the sole carbon source (*n*=3). b) The rate of ATP production measured using the ATP/ADP approach from *R. rhodochrous* incubated with a polymer film (*n*=3). Bacteria grown with no carbon source (open diamonds) died quickly as any residual carbon source was exhausted. There was no evidence of bacterial growth on LDPE (solid circles). No ATP was detected in the initial samples suggesting that the bacteria adhere immediately to the LDPE surface and were not removed sufficiently by the assay buffer. c) Measuring growth by monitoring at 600 nm of *R. rhodochrous* with no carbon source (open diamonds), on LDPE (red circles) or glucose (cyan triangles) (*n*=3). d) Measuring growth using the AlamarBlue™ assay of *R. rhodochrous* with no carbon source (open diamonds), on LDPE (red circles) or glucose (cyan triangles) (*n*=3).

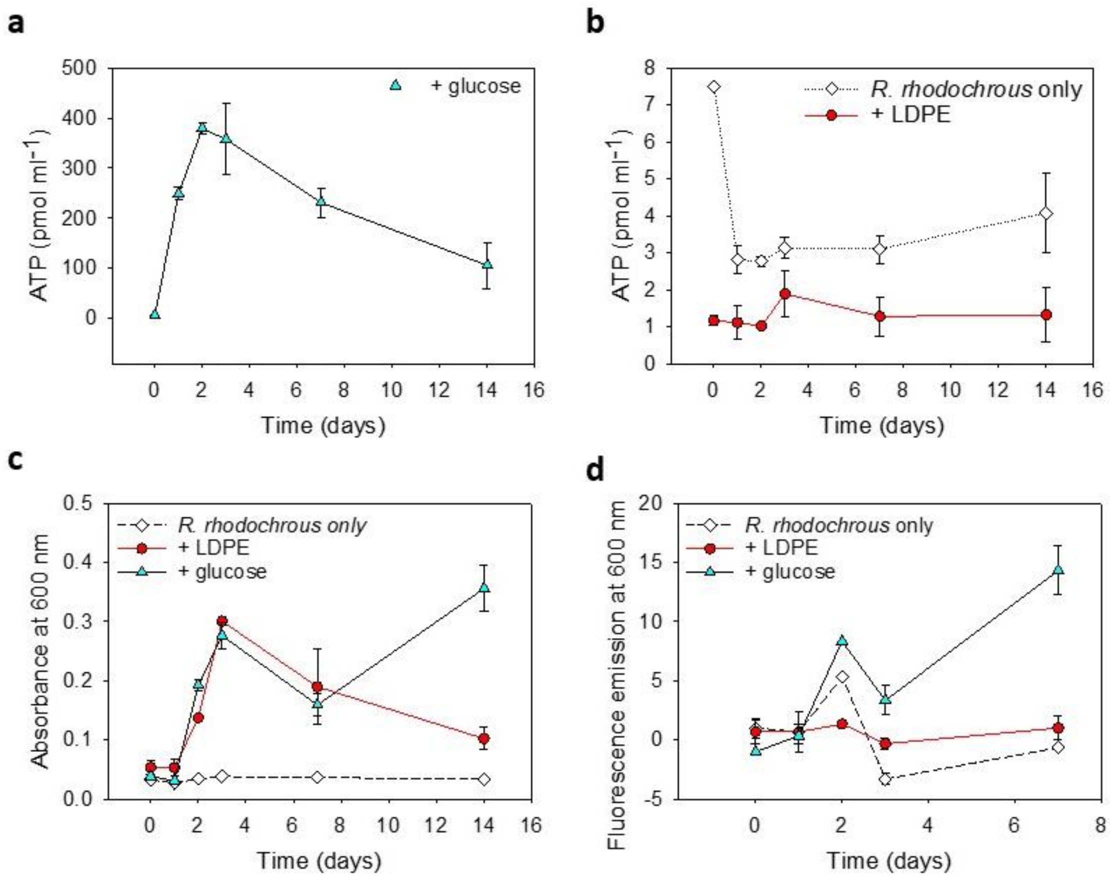

**Figure S3. Longer UV exposure increases the biodegradability of LDPE but not oxo-LDPE.**
$CO_2$ production after 35 days for *R. rhodochrous* grown on LDPE (red) or oxo-LDPE (blue) that had been exposed to UV for an increasing length of time (*n*=5, error bars represent standard error). Note how the degradation of LDPE continues to increase with prolonged exposure to UV whereas the degradation of oxo-LDPE peaks at 450 hours of irradiation.

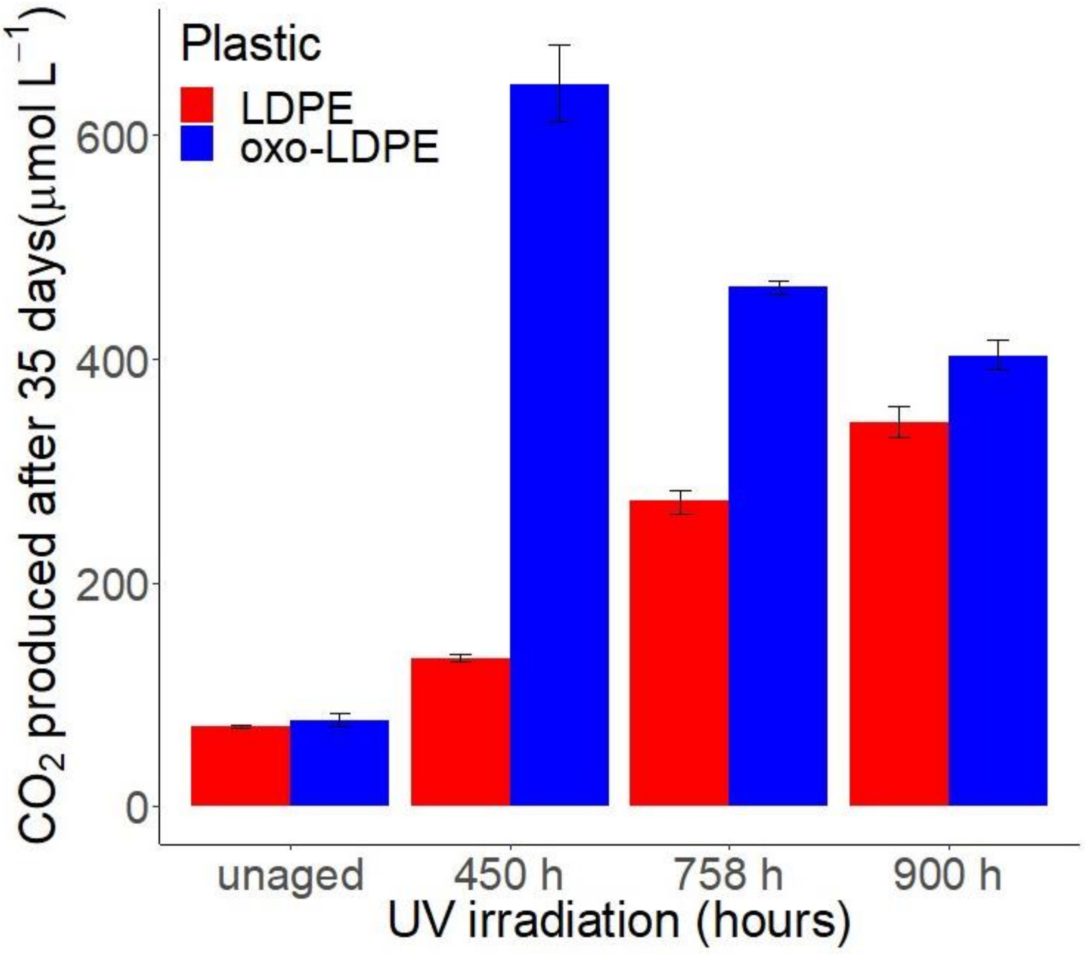

**Figure S4. The approach is highly effective at monitoring compostable plastic degradation.** *R. rhodochrous* grown with no carbon source (open diamonds/dotted line) or with a sample of a commercially available compostable plastic (yellow star/solid line) and the $CO_2$ measured over time ($n$=3, error bars represent 1 standard error).

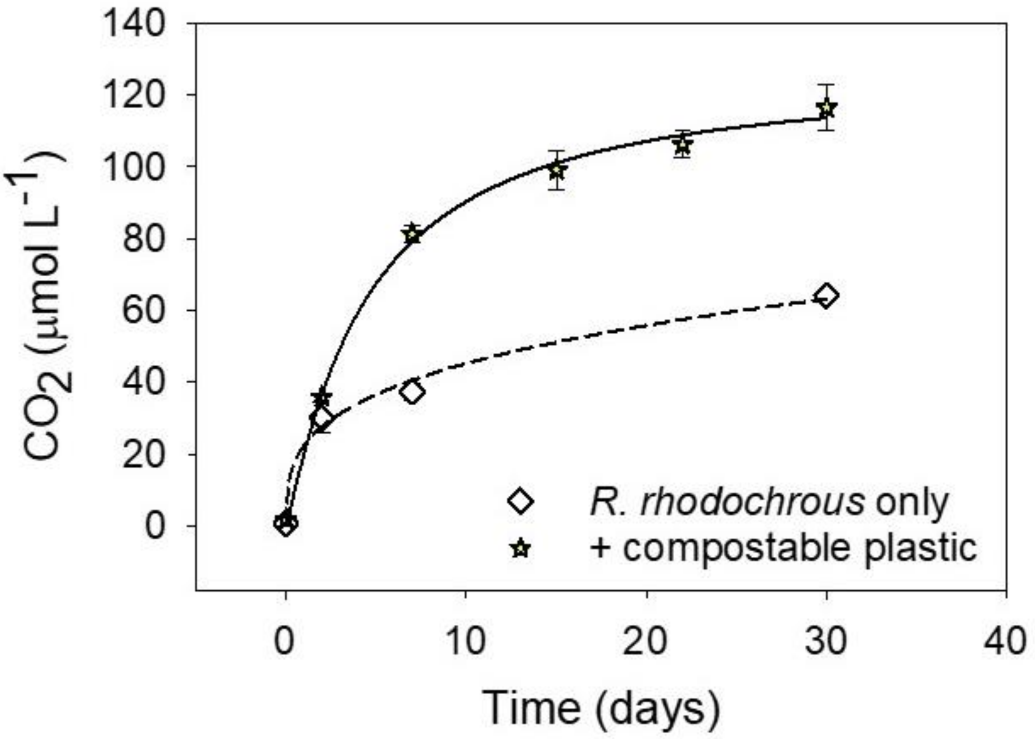
